# Mushroom body specific transcriptome analysis reveals dynamic regulation of learning and memory genes after acquisition of long-term courtship memory in *Drosophila*

**DOI:** 10.1101/364745

**Authors:** Spencer G. Jones, Kevin C.J. Nixon, Jamie M. Kramer

## Abstract

The formation and recall of long-term memory (LTM) requires neuron activity-induced gene expression. Transcriptome analysis has been used to identify genes that have altered expression after memory acquisition, however, we still have an incomplete picture of the transcriptional changes that are required for LTM formation. The complex spatial and temporal dynamics of memory formation creates significant challenges in defining memory-relevant gene expression changes. The mushroom body (MB) is a signaling hub in the insect brain that integrates sensory information to form memories. Here, we performed transcriptome analysis in the *Drosophila* MB at two time points after the acquisition of LTM: 1 hour and 24 hours. The MB transcriptome was compared to biologically paired whole head (WH) transcriptomes. In both, we identified more transcriptional changes 1 hour after memory acquisition (WH = 322, MB = 302) than at 24 hours (WH = 23, MB = 20). WH samples showed downregulation of developmental genes and upregulation of sensory response genes. In contrast, MB samples showed vastly different gene expression changes affecting biological processes that are specifically related to LTM. MB-downregulated genes were highly enriched for metabolic function, consistent with the MB-specific energy influx that occurs during LTM formation. MB-upregulated genes were highly enriched for known learning and memory processes, including calcium-mediated neurotransmitter release and cAMP signalling. The neuron activity inducible genes *hr38* and *sr* were also specifically induced in the MB. These results highlight the importance of sampling time and cell type in capturing biologically relevant transcriptional changes involved in learning and memory. Our data suggests that MB cells transiently upregulate known memory-related pathways after
memory acquisition and provides a critical frame of reference for further investigation into the role of MB-specific gene regulation in memory.

## Introduction

Learning and memory can be measured in experimental organisms by observing altered behaviour in response to manipulated experiences. The duration of behavioural changes induced by different learning and memory paradigms may be transient or stable ^1,2^. While the formation of both short-term and long-term memories require similar underlying molecular mechanisms such as calcium- and cAMP-dependent signaling pathways, only long-term memory (LTM) requires gene transcription and *de novo* protein synthesis ^3–5^. While many genes have been implicated in LTM formation ^6^, we still know very little about the spatial and temporal dynamics of gene regulation that are required for LTM.

The fruit fly, *Drosophila melanogaster*, has been a powerful model for the discovery of genes and molecular mechanisms underlying learning and memory ^5,7,8^. Transcriptome analysis has been used to identify genes expression changes in flies after the acquisition of LTM ^9–13^. Several studies have profiled transcriptional changes in whole heads ^9–11^ which has led to the identification of genes that are required for LTM ^9,11^. Despite the success of these whole head studies, it is clear that LTM requires only a subset of neurons that are both spatially and temporally regulated ^14–16^. Cell-type specific analysis of different neuronal subsets will be required to identify gene expression changes that are critical for memory ^17^.

The mushroom body (MB) is a region of the fly brain that is critical for normal memory ^18,19^. This synaptically dense structure appears as a pair of neuropils each consisting of ~2000 neurons with three distinct neuronal subtypes (α/β, α’/β’, and γ) that contribute the formation of 5 distinct lobes α, α’, β, β’, and γ ^20^. Intrinsic MB neurons, called Kenyon cells (KC), form a hub for integration of sensory information from over 200 olfactory projection neurons and 20 different modulatory dopaminergic neurons ^21^. Sensory information is processed in the MB and relayed through just 21 MB output neurons (MBONs) ^22^. Because of its essential role in memory, the MB is a logical starting point in the search for LTM-dependent gene expression changes. MB-specific transcriptome analysis has led to the discovery of additional genes that are important for LTM, however, similar to whole head analysis, no consistent LTM-dependent gene regulatory patterns have been observed ^12,13^. This lack of consistency - in both whole head and MB-targeted transcriptome analysis - is likely due to a range of factors including differences in sampling time, e.g. 30 minutes vs. 12 hours after memory acquisition. Indeed, gene expression changes are known to vary at different time points after memory acquisition ^23^. It is also likely that memory dependent gene expression changes will differ based on physiological differences resulting from the different memory paradigms used, e.g. appetitive vs. aversive olfactory conditioning. Indeed, it has been shown that neuron activity regulated gene expression is highly specific not only to neuron type but also to the stimulation paradigm ^24^. Therefore, in order to identify gene regulatory mechanisms that are essential for memory, it will be important to investigate different memory paradigms, different neuronal subsets, and different time-points during and after memory acquisition.

Courtship conditioning is a well-established learning and memory paradigm that has been commonly used to investigate the molecular mechanisms underlying memory ^19,25–27^. Courtship conditioning relies on male courtship behaviour being modifiable in response to sexual rejection from a mated unreceptive female ^28,29^. After experiencing sexual rejection males show reduce courting attempts with other pre-mated females; an effect which can persist for several days ^19,27^. Courtship memory forms via an enhanced behavioural response to the pheromone cis-vaccenyl-acetate (cVA), which is deposited on females by males during prior mating attempts ^30^. The MB is required for the acquisition of normal long-term courtship memory ^19^. While courtship conditioning has molecular properties similar to other memory paradigms ^31^, it is distinct in that it manipulates a complex, naturally occurring behaviour with minimal experimental interference ^30–32^. This makes courtship conditioning an attractive model that takes advantage of a robust but ethological form of memory.

Here, we use INTACT (isolation of nuclei tagged in a specific cell type)^33^ to profile gene expression in MBs at two time points after the acquisition of long-term courtship memory. We find a dynamic effect on the regulation of learning and memory genes during LTM formation in MBs. Many known learning and memory genes are transiently upregulated in MBs one-hour after memory acquisition and return to baseline levels after 24 hours. This effect is specific to MBs, as whole head transcriptome analysis did not reveal gene regulatory changes in known memory associated biological pathways. This suggests a high demand for classic learning and memory genes in MBs after the acquisition of courtship memory and highlights the importance of sampling time and cell type in the detection of biologically relevant transcriptional changes underlying memory.

## Results

### MB-unc84 males display normal long-term courtship memory

The aim of this study was to identify MB-specific transcriptional changes that occur after the acquisition of long-term courtship memory. To achieve this, we used INTACT ^33–35^ to isolate MB nuclei from fly heads, 1 h and 24 h after courtship conditioning (**Figure 1A**). We adapted a previously described INTACT protocol that employed a *UAS*-*unc84::GFP* transgene ^33^. Unc-84 is a *Caenorhabditis elegans* nuclear envelope protein, and when coupled with GFP, Unc84::GFP labeled nuclei can be immunoprecipitated from nuclear extracts derived from frozen tissue using an anti-GFP antibody. To drive expression of *UAS*-*unc84::GFP* in the MB, we used the *R14H06*-*GAL4* driver line from the Janelia flylight collection ^36^. This driver is highly specific for the α/β and γ neurons of the mushroom body (**Figure 1A**). We generated flies that were heterozygous for both the *UAS*-*unc84* transgene and the *R14H06*-*GAL4* driver, which are hereafter referred to as *MB*-*unc84*.

**Figure 1.**
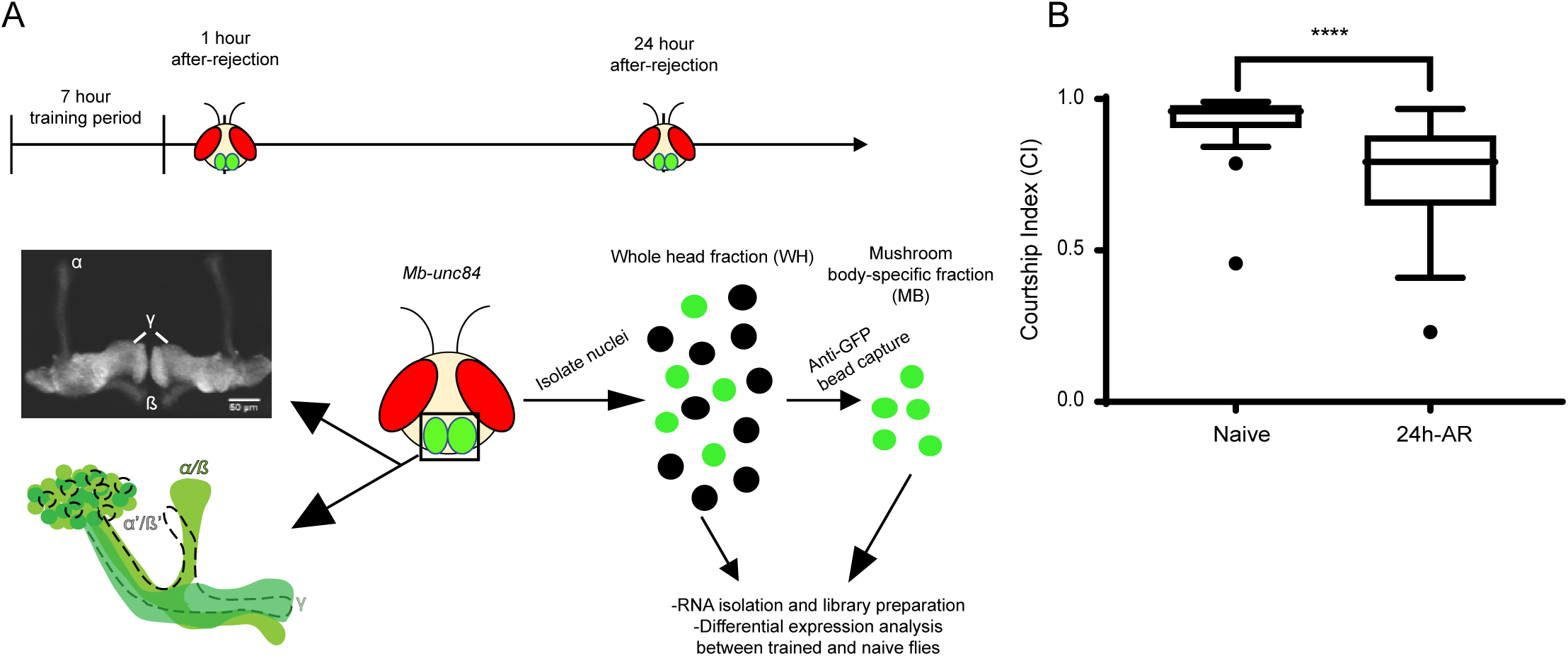
Schematic of the experimental design and validation of courtship conditioning to induce LTM. **A)** Long-term memory (LTM) was induced in flies using a previously established seven-hour courtship conditioning protocol. Following training, flies were collected for downstream transcriptome analysis at two time points: one hour and 24 hours after-rejection. Flies used for analysis were heterozygous for both *R14H06*-*GAL4* and a GFP-bound nuclear membrane tag *UAS*_*unc84*-*2XGFP* (unc84), referred to as *Mb*-*unc84.* The *R14H06* GAL4 line was used to drive the expression of *unc84* as it is specific to the Kenyon cells of the α/β and γ lobes of the mushroom body, regions still requiring further investigation during LTM. Fly heads were obtained from samples of 50-60 flies through flash freezing with liquid nitrogen, followed by vortexing and separation through a series of sieves. Fly heads were then suspended in homogenization buffer and nuclei released into solution through chemical and physical agitation of the cell membrane. A whole head fraction (WH) was then taken from this homogenate as a representation of the whole fly head, containing both MB-specific GFP nuclei and untagged non-MB nuclei. Anti-GFP bound beads were then used to isolate GFP-positive nuclei from the whole head homogenate to represent the mushroom body-specific fraction (MB). RNA was then isolated from both WH and MB-fractions, cDNA libraries prepared and next-generation sequencing performed. Differential expression (DE) analysis was then performed between trained flies and untrained, naïve flies. **B)** To provide evidence of normal LTM functioning in flies used for analysis, a subset of flies from each day of training were tested for retained courtship suppression 24 hours later (24h-AR). Individual trained and naïve male flies were introduced to a pre-mated female, videoing their interactions for 10 minutes and quantifying observed courtship behaviours. The amount of time spent courting is represented as a courtship index (C.I.). For this experiment, n = 23 and n =29, respectively, for naïve and trained flies; ^∗∗∗∗^ p < 0.001 Mann-Whitney U-test.

To induce long-term courtship memory, *MB*-*unc84* males were paired with an unreceptive mated female for seven hours. Flies for transcriptome analysis were flash frozen at 1 h and 24 h after this period of sexual rejection (1h-AR and 24h-AR, **Figure 1A**). These time points were selected to capture both early and late stages after memory acquisition. We avoided sampling during the rejection period to avoid the direct effect of being paired with a female ^37,38^. A minimum of four biological replicates was obtained for each time point. In parallel with these collections, we tested a subset of *MB*-*unc84* flies to confirm the induction of normal long-term courtship memory. Indeed, at 24h-AR *MB*-*unc84* males showed a robust reduction in courtship behaviour in comparison to naïve males (**Figure 1B**; p < 0.001 Mann-Whitney *U*-test). This observed courtship suppression in *MB*-*unc84* flies was in line with expected values from the literature ^19,25,27^, demonstrating that *UAS*-*unc84::GFP* expression in the MB does not interfere with normal courtship memory.

### INTACT yields high-quality MB-enriched RNA

To provide evidence that our approach could obtain nuclei specific to the MB, we used fluorescent microscopy to measure the proportion of GFP-positive nuclei present in whole head (WH) extracts, compared to INTACT MB samples. WH nuclear extracts obtained from *MB*-*unc84* flies contained 8% GFP positive nuclei (**Figure 2A**). Note that this is likely an overestimation, as we only analyzed fields of view containing GFP-positive nuclei, which were not present throughout the slide. After immunoprecipitation of nuclei from WH extracts using anti-GFP bound beads, about 90% of nuclei were GFP-positive, indicating a high level of specificity of our INTACT protocol (**Figure 2A**).

**Figure 2.**
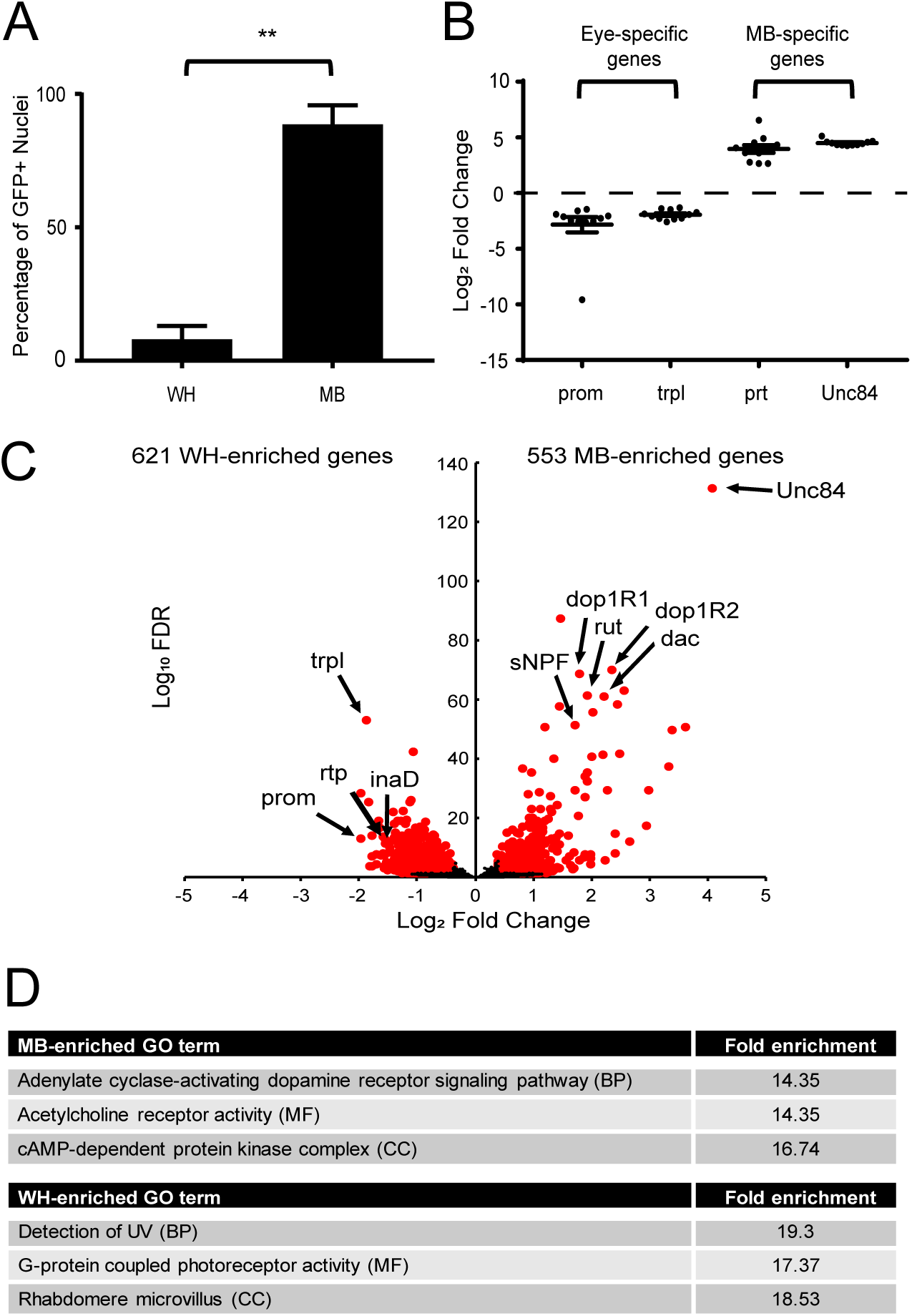
INTACT yields high-quality MB-enriched RNA. **A)** Average percentage (± SD) of GFP positive nuclei in WH and MB samples (n=3) obtained using INTACT. MB samples show a 13.8-fold increase (^∗∗^ p - < 0.01, Student’s t-test) in the percentage of GFP positive nuclei compared to WH samples. **B)** Scatter dot plot of log2 fold changes for a selection of genes with known expression in the eye (*prom*, *trpl*), representing the whole fly head, the mushroom body (MB) (*prt*), as well as the genetically expressed nuclear tag *unc84.* Fold changes were calculated for biologically paired WH and MB samples (n=11) using normalized counts across all samples. Error bars represent standard error of the mean. **C)** Differential expression (DE) analysis was performed on all MB and WH samples after removing genes with less than 50 mean normalized counts. Volcano plot shows the results of this analysis, which revealed 553 and 621 DE genes (q < 0.05, fold difference > 1.3), respectively, in MB-enriched and WH-enriched samples. A selection of genes previously known to be enriched in both MB-cells (*rut*, *dac*), KC’s (*sNPF*, *Dop1R1*, *Dop1R2*) and WH-cells (*trpl*, *prom*, *rtp*, *inaD*), as well as *unc84*, are highlighted. **D)** Gene ontology (GO) analysis was performed separately on lists of MB-enriched and WH-enriched DE genes. Significant terms (p < 0.05, Fisher Exact with FDR multiple test correction, minimum four genes) with the highest enrichment for biological processes (BP), molecular functions (MF) and cellular components (CC) are displayed for both MB and WH-enriched DE lists.

Next, INTACT was used to extract MB nuclei from *MB*-*unc84* flies’ heads at 1h-AR and 24h-AR, as well as from naïve flies matched for age and time-of-day. For each MB sample, we also obtained RNA from nuclei present in the biologically paired WH input for comparison. After verification of RNA quality, sequencing libraries were prepared from both WH and MB samples. Completed libraries were sequenced and reads were aligned to the *D. melanogaster*genome. Samples that had >10 million genic reads were included for downstream analysis, resulting in a total of 12 MB samples: four naïve, four 1h-AR, four 24h-AR - and 12 WH samples - five naïve, three 1 h-AR, four 24h-AR (**Table S1**).

To confirm consistent MB enrichment in MB samples we examined gene expression differences between WH and MB samples. DESeq2 was used to normalize gene counts between all MB and WH samples and genes with less than 50 mean normalized counts across all samples were removed, leaving a total of 11941 genes with sufficient coverage. Log_2_ fold changes were then calculated for biologically paired WH and MB samples. As expected, eye-specific genes like *prom* and *trpl* were underrepresented in MB-samples, while MB-enriched genes, such as *portabella* and *unc84* were overrepresented in the MB samples (**Figure 2B**). Notably, *unc84* expression was highly enriched and highly consistent across all biological replicates suggesting a high degree of consistency in MB-enrichment after INTACT.

To provide further evidence that the nuclei we isolated displayed MB-specific gene expression profiles we performed differential expression analysis between all MB and WH samples. We identified 553 and 621 genes (FDR < 0.05, fold difference > 1.3) that were significantly enriched in either MB or WH samples, respectively (**Figure 2C**; complete list in **Table S2**). Many known MB-expressed genes, including *rutabaga*, *dunce*, *prt*, *eyeless*, *twin of eyeless*, and *dachshund* were among the most differentially expressed MB-enriched genes ^7,39–41^. In contrast, several eye-specific genes, such as *prom*, *trpl*, *inaD*, and *rtp*, were among the most differentially expressed WH-enriched genes (**Figure 2C**). Additionally, we compared MB-enriched genes to cell surface receptors that were found to be characteristically expressed in α/β and γ KC’s when compared to MBONs ^12^. Indeed, many of these receptors were also found to be enriched in our dataset including: *Dop1R2*, *Dop1R1*, *Dop2R*, *5*-*HT1B*, *Oamb*, *OctβR*, *sNPF*, *GluRIB*, *Ir68a*, *CCKLR*-*17D1*, *CCKLR*-*17D3*, *GluRIB*, *and mAChR*-*A* (**Table S2**). Finally, we examined gene ontology (GO) terms enriched for biological processes (BP), molecular functions (MF), as well as cellular components (CC), among our lists of MB-enriched and WH-enriched genes (**Figure 2D**, **Table S3**). The most enriched GO terms for MB-enriched genes were “cAMP-dependent protein kinase complex” (CC) and “adenylate cyclase-activating dopamine receptor signaling pathway” (BP) (**Figure 2D**), which fits with the known importance of dopaminergic modulatory neurons and cAMP signaling in memory formation in the MB ^22,30,42,43^. The most enriched GO term for MF was “acetylcholine receptor activity”, consistent with previous studies which showed that MB KC’s are cholinergic and receive input from cholinergic olfactory projection neurons ^12,44,45^. In contrast, the most enriched GO terms for the WH enriched genes were all related to eye function, including “detection of UV” (BP), “G-protein coupled photoreceptor activity” (MF) and “rhabdomere microvillus” (CC) (**Figure 2D**). Taken together, analysis of genes that are differentially expressed between WH and MB samples revealed a pattern of gene expression that is highly consistent with an effective MB enrichment.

### Gene expression changes after memory acquisition

Next, we used DESeq2 to identify genes that were differentially expressed (DE) in response to courtship conditioning by comparing 1h-AR and 24h-AR to naïve flies. For both WH and MB samples we observed more DE genes at 1 h-AR than at 24h-AR (n=322/23, n=302/20, for WH and MB sample 1h-AR/24h-AR, respectively). There was some overlap in DE genes between 1h-AR and 24h-AR, and between WH and MB samples, however, most DE genes identified in WH and MB samples were different (**Figure 3A**). To investigate trends in gene expression after courtship conditioning we compiled a list of all DE genes that were differentially expressed in at least one of the three pairwise comparisons: 1h-AR vs. naïve, 24h-AR vs. naïve, and 1h-AR vs. 24h-AR (**Table S4**). This led to the identification of 332 and 342 DE genes for WH and MB samples, respectively. For each tissue, we performed *k*-means clustering on log2 fold changes at 1h-AR and 24h-AR (**Figure 3B and 3C, Table S5**). In WH samples, four clusters were identified with two distinct trends: cluster 1 and 2 (n=22 and 127) contained genes that were downregulated at 1h-AR and either reduced or not changed at 24h-AR (WH-down, **Figure 3B and Table S5**). Cluster 3 and 4 (n = 72 and 114) contained genes that were upregulated at 1h-AR and either less upregulated or not changed at 24h-AR (WH-up - **Figure 3B and Table S5**). For MB samples *k*-means clustering revealed five clusters with three distinct expression trends. Cluster 1 (n = 30) contained genes that were downregulated at 1h-AR and upregulated 24h-AR. Clusters 2, 3, and 4 (n=2, 13 and 120, respectively) contained genes that were downregulated at 1h-AR and either downregulated or not changed at 24h-AR (MB-down - **Figure 3C and Table S5**). Cluster 5 (n = 174) contained genes that were upregulated at 1h-AR and either upregulated or not changed at 24h-AR (MB-up- **Figure 3C and Table S5**). This clustering allowed us to identify gene groups with similar expression trends and emphasized the relatively strong effect of sexual rejection at 1h-AR.

**Figure 3.**
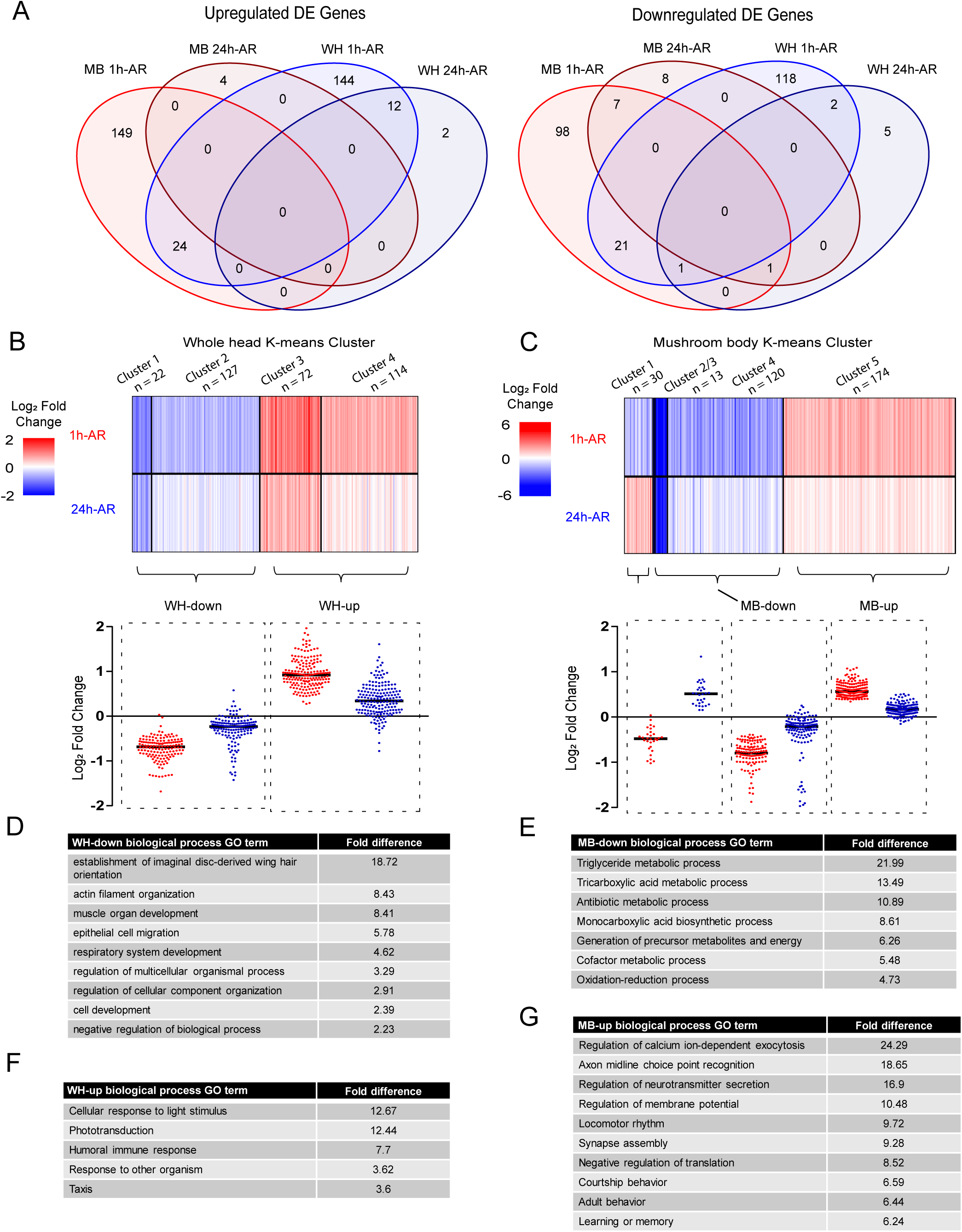
Differential expression and clustering analysis of MB and WH sequencing results. **A)** Venn diagram shows overlap between MB and WH differentially expressed (DE) gene lists (q < 0.05, fold difference 1.3 up or down) for both upregulated and downregulated genes. **B)** A list of 336 WH DE genes (q < 0.05, fold difference 1.3 up or down) was determined through DE analysis of all three experimental conditions (1h-AR vs naïve, 24h-AR vs naïve and 24h-AR vs 1h-AR). Log2 fold change data was obtained for significant genes at both one hour, as well as 24-hour time points and clustered using *k*-means. Four clusters were identified with two distinct trends. Cluster 1 and 2 were downregulated at both time-points (WH-down) and cluster 3 and 4 were upregulated at both time-points (WH-up). Heatmap shows the individual log2 fold changes for each clustered gene. Scatter dot plot shows log2 fold changes for genes with similar expression trends. **C)** A list of 343 MB DE genes (q < 0.05, fold difference 1.3 up or down) was determined using the same approach as that used for WH DE genes. Log2 fold change data was obtained for significant genes at both one hour, as well as 24-hour time points and clustered using k-means. Five clusters were identified with three distinct trends. Cluster 1 was downregulated 1h-AR and upregulated 24h-AR. Cluster 2, 3 and 4 were downregulated at both time-points (MB-down). Cluster 5 was upregulated at both time-points (MB-up). Heatmap shows the individual log2 fold changes for each clustered gene. Scatter dot plot shows log2 fold changes for genes with similar expression trends. **D)** GO analysis results using PANTHER for biological processes for WH-down genes (p <0.05, Binomial test with Bonferroni correction, sorted by hierarchical view). The top nine GO terms, representing each ontology class and containing at least five genes, are displayed, sorted by fold enrichment. **E)** GO analysis results using PANTHER for biological processes for MB-down genes (p <0.05, Binomial test with Bonferroni correction, sorted by hierarchical view). The top seven GO terms, representing each ontology class and containing at least five genes, are displayed, sorted by fold enrichment. **F)** GO analysis results using PANTHER for biological processes for WH-up (p <0.05, Binomial test with Bonferroni correction, sorted by hierarchical view). The top five GO terms, representing each ontology class and containing at least five genes, are displayed, sorted by fold enrichment. **G)** GO analysis results using PANTHER for biological processes for MB-up genes (p <0.05, Binomial test with Bonferroni correction, sorted by hierarchical view). The top 10 GO terms, representing each ontology class and containing at least five genes, are displayed, sorted by fold enrichment.

### Courtship conditioning is associated with MB-specific downregulation of metabolic genes

To investigate the functions of genes that are differentially expressed in response to courtship conditioning, we first performed GO enrichment analysis for gene clusters with similar expression trends. For WH-down genes (n = 149, **Figure 3B**) we observed, almost exclusively, enrichment of GO terms related to development, for example, “metamorphosis”, “cell differentiation”, and “cell migration” (**Figure 3D and Table S6**). For MB-down genes (n = 135, **Figure 3C**) we observed enrichment only of GO terms related to metabolism (**Figure 3E and Table S6**). In fact, over half (n=73) of the MB-specific downregulated genes are annotated with the term “metabolic processes” (**Table S6**). Notably, there was no overlap in enriched GO terms between WH-down and MB-down genes. The highly specific effect of courtship conditioning on the regulation of metabolic genes in the MB is very interesting as it has been shown that increased energy metabolism in the MB is required for formation of olfactory LTM ^46^. Consistent with our observations, the energy influx observed in the MB during LTM formation is not seen in other brain regions ^46^. Thus, downregulation of many metabolic genes in the MB at 1h-AR may reflect the shifting metabolic state in the MB that is required for memory formation.

### Courtship conditioning is associated with MB-specific upregulation of synaptic proteins and learning and memory pathways

For WH-up genes (n= 186, **Figure 3B**) all enriched GO terms were related to biological responses, such as “cellular response to light stimulus”, “humoral immune response”, “response to other organism”, and “taxis” (**Figure 3F and Table S6**). Indeed, 62 of the WH-up genes were annotated with the term “response to stimulus” (**Table S6**). GO terms related to biological response were also enriched for MB-up genes (n = 174, **Figure 3C**). There were 5 enriched GO terms common to WH-up and MB-up genes (“response to light stimulus”, “response to abiotic stimulus”, “response to stimulus”, “response to external stimulus”, and “taxis”) (**Table S6**). Yet the MB-up gene group showed many more enriched GO terms - 181 compared to 15 for WH-up - suggesting a high level of functional relatedness in this gene group. Using annotated protein-protein and genetic interactions, we identified a network 54 MB-up genes (**Figure 4**). This network was comprised of genes encoding ion channels, transcription factors, RNA binding proteins, and genes with functional annotations related to synapse formation, synaptic signaling, behaviour, and learning and memory (**Figure 4**). Interestingly, the most enriched GO categories that were unique for MB-up genes were related to synaptic plasticity (e.g. “regulation of calcium ion-dependent exocytosis”), behaviour (e.g. “courtship behaviour”), and memory (“e.g. “learning or memory”) (**Figure 3G**). Taken together, these results suggest that MB-up genes encode a highly interactive group of proteins with biological relevance to learning and memory.

**Figure 4.**
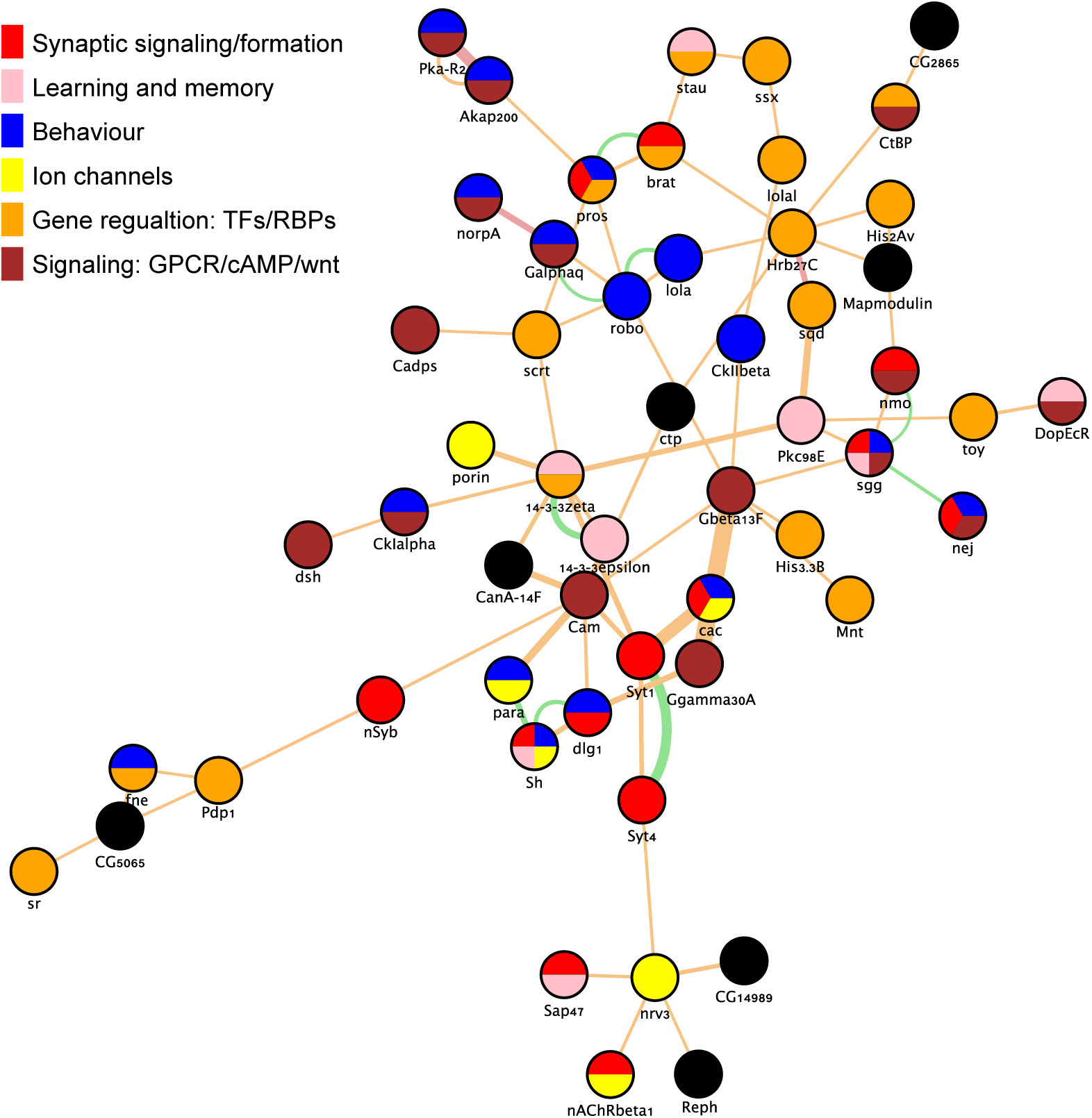
Network analysis of genes that are upregulated in the MB in response to courtship conditioning. Of 178 genes in the MB-up group, 54 form a single network based on a subset protein-protein and genetic interactions that are annotated in geneMANIA (see methods). Each node is colour coded to represent selected

Next, we manually curated the MB-up gene group to illustrate how they may be represented in memory-relevant molecular pathways in MB KCs (**Figure 5**). During learning and memory formation KCs receive olfactory input from over 200 olfactory projection neurons (PNs) that synapse with the dendrites of the calyx ^47^. Olfactory signals are reinforced to form memories by sensory signals from modulatory dopaminergic neurons, which synapse at discrete locations along the axons of the MB lobes ^22^. In courtship conditioning, the primary olfactory signal is thought to be the pheromone cVA which is deposited on females by males during mating ^48^. Courtship memory is formed when cVA is paired with sexual rejection, which is conveyed to the MB gamma lobe via a specific class of dopaminergic neurons ^30^. Long-term courtship memory is also dependent on the production of the hormone ecdysone, which also can also act as an input signal to KCs ^49,50^. Olfactory PNs are cholinergic and are thought to stimulate KCs through activation of nicotinic acetylcholine receptors (nAChRs), which are ligand-gated channels that induce calcium influx into KCs. Calcium influx is required for downstream signaling associated with synaptic plasticity and memory formation. Among MB-up genes, we noted several genes involved in receiving olfactory signals and mediating downstream calcium dependent signaling (**Figure 5**). These included genes encoding three nAChR subunits (*nAChRα1*, *nAChRα6*, *nAChRβ1*), the acetylcholinesterase (*ace*) involved in acetylcholine recycling, the voltage-gated calcium channel *Ca*-*beta*, the calcium-activated signalling proteins PLC and PKC, the and the calcium-binding messenger calmodulin (cam). Many MB-up genes also encode proteins involved in receiving modulatory signals, and in the cAMP signaling pathway that is activated by these signals during memory formation (**Figure 5**). Notably, we identified MB-specific upregulation of four G-protein coupled receptors (GPCR). These included oamb, hec, and SIFaR, all known to be involved in male courtship behaviour ^51–53^, and DopEcR, an atypical GPCR that responds to both dopamine and ecdysone, and is essential for cAMP signal activation during courtship memory ^50^. We identified five MB-up genes encoding components of the heterotrimeric G-protein complex (Gαq, Gβ13F, Gγ30A, Gαo, Gγ1) which act directly downstream GPCRs to induce adenylate cyclase activity and production of cAMP. Several downstream cAMP signaling components were also upregulated specifically in the MB, including regulatory subunits of protein kinase A (PKA-R2), the PKA anchoring protein (Akap200), cAMP-gated ion channels (*Ih*, *Cngl*), and the CREB-binding protein, nej, a histone acetyltransferase that is thought to be involved in LTM-associated gene expression ^54,55^. Thus, many MB-up genes are directly related to receiving and processing the signals that induce courtship memory.

**Figure 5:**
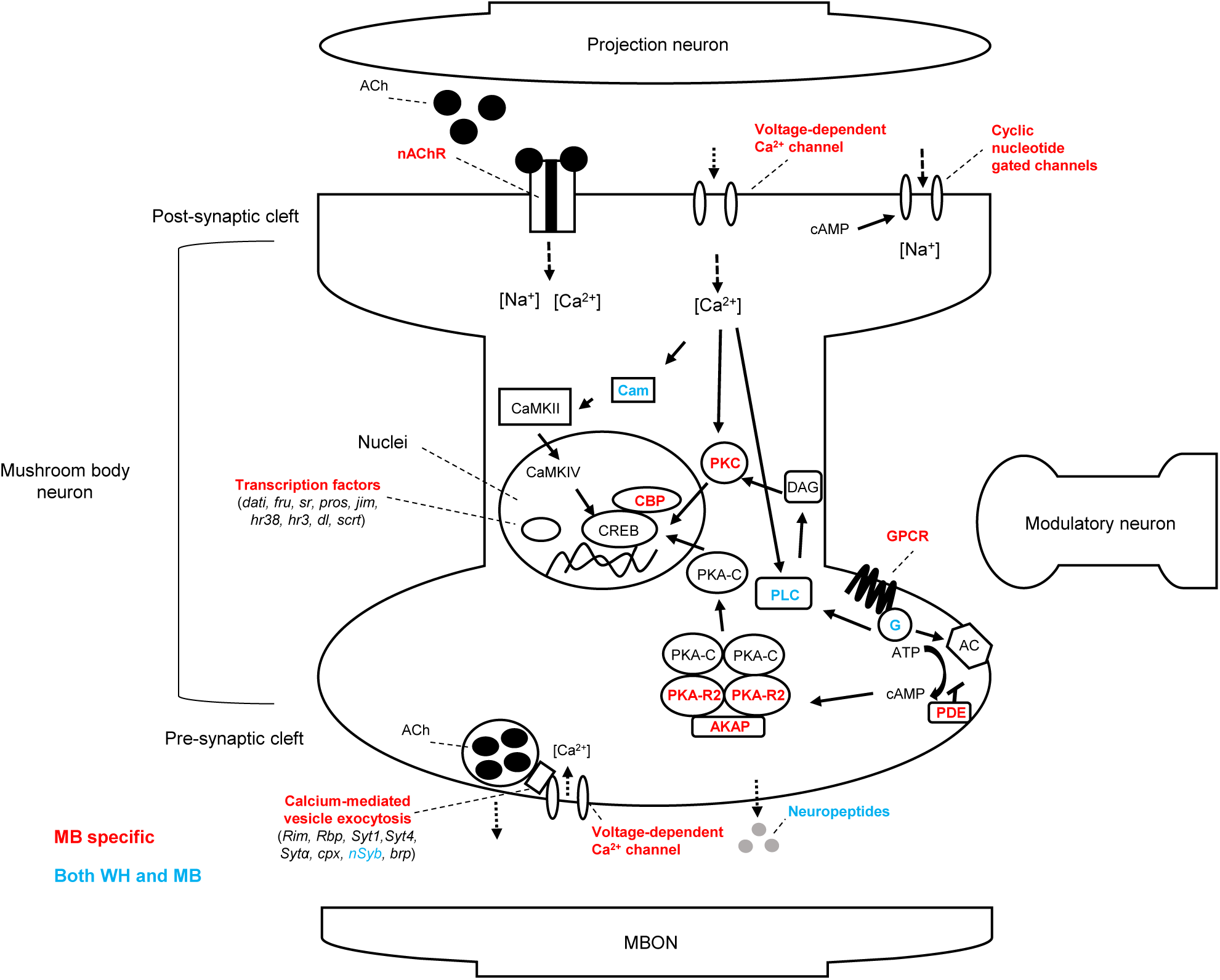
Schematic representation of molecular pathways underlying memory in the mushroom body. Manually curated diagram of memory-relevant molecular pathways in MB Kenyon cells which were identified to be differentially expressed in the MB-up gene group (shown in red). Calcium-dependent, cholinergic and cAMP signaling pathways are among the molecular pathways represented. Additionally, genes encoding proteins involved in calcium in calcium-mediated presynaptic neurotransmitter release, as well as differentially expressed transcription factors are shown.

KC axons provide presynaptic output to 21 MBONs. Several MB-up genes encode proteins involved in calcium-mediated presynaptic neurotransmitter release, including the synaptic vesicle docking proteins RIM and RBP, the synaptotagmins (Syt1, Syt4, Sytα), components of the SNARE complex (cpx and nSyb), the presynaptic calcium channel cacophony, and the active zone marker brp (**Figure 5**). We also observed upregulation of two neuropeptides, Nplp2, and sNPF. sNPF is has been shown to act synergistically with ACh in communicating to MBONs in the context of olfactory memory formation ^44^. Thus, many MB-up genes are involved in transmitting memory signals to MBONs (**Figure 5**).

Finally, we also observed upregulation of many genes encoding transcription factors and RNA binding proteins. RNA binding proteins like stau and Orb2 are thought to be involved in LTM formation through localized regulation of translation at synapses ^9,56^. Some of the transcription factors in the MB-up group have known roles in courtship behaviour, such as dati, fru and pros^57–59^. Interestingly, we identified MB-specific upregulation of sr and HR38, which are transcription factors that have been proposed as markers of neuron activation in insects ^24,60,61^.

## Discussion

Understanding transcriptional changes that are required in neurons to mediate LTM is an important challenge in neuroscience. Many studies have identified gene expression changes after memory acquisition in *Drosophila* ^9–12^ and this approach has been used to identify new genes involved in memory formation ^9,11,12^. However, we still understand very little about the spatial and temporal requirement for transcription in LTM. When are critical memory genes activated and in which neurons? Here, we used MB-specific transcriptional profiling to identify gene expression changes that occur in response to courtship conditioning, an ethological memory paradigm that is commonly used in *Drosophila.* This analysis revealed gene expression changes in established learning and memory pathways that occurred for the most part at 1 hour after courtship rejection, but not after 24 hours. Importantly, memory related pathways were only differentially regulated in the MB and not in biologically paired WH samples. These results suggest that memory related biological processes are transiently upregulated in the MB after memory acquisition and illustrate the importance of sampling time and cell type in the identification of biologically relevant gene regulation in LTM.

In our study, we compared males that experienced sexual rejection to naïve socially isolated males. Although samples were collected at least one hour after exposure to a female fly, it is impossible to conclusively differentiate between transcriptional changes that occur because of sexual rejection - and long-term memory formation - and changes that might happen in response to any social interaction. Previous studies have looked at gene expression changes that occur in whole heads in response to courtship, male-male interactions, and mating ^37,38^. As could be expected, in WH samples we do observe a significant overlap with those studies (36 genes, 1.5-fold enrichment, p = 0.009). This suggests that some gene expression changes in whole heads represent general responses to social interactions. In contrast, we see no significant overlap between genes identified in these studies and genes that we observe to be changed in the MB. In addition, the MB is not required for normal male-female interactions like courtship behaviour or mating as MB-ablated flies reproduce normally and even show normal learning in response to sexual rejection ^18,19^. Therefore, it is reasonable to suggest that MB-specific gene expression changes we observed are likely related to memory acquisition.

The nature of the genes and biological pathways that are differentially expressed specifically in the MB after memory acquisition strongly suggest a role in memory. The MB-specific changes in metabolic gene expression that we observe correlate well with the MB-specific energy influx previously described for appetitive olfactory conditioning ^46^. MB-specific energy consumption during LTM is likely mediated by post-translational mechanisms such as the phosphorylation of the pyruvate dehydrogenase complex - not transcription. However, such a dramatic metabolic shift is very likely to indirectly affect the expression of metabolic genes, which could explain why more than half of MB-down genes identified in our study are linked to metabolism.

Genes that are upregulated in the MB after memory acquisition show a remarkable correlation with known memory pathways. From post-synaptic receptors, to signaling pathways, to presynaptic neurotransmitter release mechanisms, nearly all known aspects of memory related synaptic plasticity are accounted for (**Figures 3-5**). Some genes, like *stau* and *fru*, were previously shown to be upregulated in whole heads during and after olfactory ^9^ or courtship memory ^10^, respectively, further validating our approach and results. However, in general, other *Drosophila* memory transcriptome studies have not observed such a profound effect on known memory related genes and pathways ^9–12^. This is likely due to both the sampling time and cell type we investigated.

Certainly, memory specific transcriptional signals would be diluted in whole head analysis ^9–11^. Widmer *et al.* used MB specific transcriptome analysis at 24 hours after appetitive olfactory memory acquisition, and consistent with our findings, identified a limited number of expression changes ^13^. Crocker *et al.* used cell-specific patch clamping to investigate gene expression from MB neurons at 30 minutes after memory acquisition, however, they identified very few differentially expressed genes, likely due to pooling of many samples that were conditioned with different odors ^12^. The fact that we observed many expected memory genes and pathways to be induced in the MB suggests that we have captured a critical time point for gene regulation in the formation of long-term courtship memory.

Many studies in mouse have profiled transcriptional changes in the hippocampus in response to fear conditioning and other memory paradigms. Consistent with our observations, these studies show more gene expression changes 30 minutes after memory acquisition, and not at later time points ^23^. In general, however, these studies do not identify widespread differential expression of classic learning and memory pathways as we do in the fly MB. In mouse, across many different studies, fear conditioning consistently invokes strong activation of immediate early genes such as cFos, which are known to be induced in response to neuron firing. In insects, neuron activity induced genes have been more elusive, however, two genes, *hr38* and *sr*, are consistently upregulated in response to a variety of neuronal activation stimuli in flies and other insects^24,60,61^. It is very interesting that we observe these two genes to be specifically activated in the MB in response to sexual rejection. No other *Drosophila* memory-related transcriptome study has identified induction of these genes ^9–13^, except for Crocker *et al.* who did identify hr38 induction in the MB α/β cells at 30 min after memory acquisition, albeit with a borderline q-value (0.058) ^12^. This suggests that our MB-specific analysis, coupled with an appropriate sampling time, has revealed a parallel mechanism to mammals that has not previously been observed in flies, where the induction of neuron activity induced genes is observed following memory acquisition.

In the future, it will be important to further refine the cell types and sampling times to fully understand transcriptional dynamics associated with memory formation. Indeed, even by focusing on less than 2000 MB cells, the actual circuit involved in the formation and long-term maintenance of the memory is likely composed of far fewer cells. The specific circuits that are required for courtship memory and other memory forms are being elucidated rapidly^16,31^ and tools are now becoming available to label these cell populations for genomic analysis ^12,33,62^. It is likely that further focus on more discrete cell populations will be required to fully understand gene activation in LTM.

## Materials and Methods

### Fly strains

All *Drosophila melanogaster* strains were cultured at 25° C and 70% humidity on a 12:12 light-dark cycle. Cultures were raised on a standard medium (cornmeal-sucrose-yeast-agar) supplemented by the mold inhibitors methyl-paraben and propanoic acid ^25^. To utilize the UAS/GAL4 expression system flies containing the MB-specific GAL4 line *R14H06*-*GAL4* (Bloomington Stock #48667) were crossed to flies with *UAS*_*unc84*-*2XGFP* (UAS-unc84::GFP), which encodes a *C. elegans*-derived nuclear tag combined with green fluorescent protein (GFP). *R14H06*-*GAL4* flies were generated by the Janelia Farm Flylight project ^36^ and obtained from Bloomington stock center and *UAS*-*unc84::GFP* flies were donated by Gilbert L. Henry, Janelia Farm Research Campus ^33^. For courtship conditioning assays and transcriptome analysis heterozygotes generated by crossing *UAS*-*unc84::GFP*; *R14H06*-*GAL4* flies to *P{CaryP}attP2* (Bloomington stock# 36303). The resulting progeny referred to as MB-unc84 have the genotype *X*; *UAS*-*unc84::GFP/*+;*R14H06*-*GAL4/attP2.* Courtship conditioning was performed using pre-mated, wild-type females with a Canton-S and Oregon-R mixed genetic background generated by J. Kramer.

### Courtship conditioning and sample collection

Long-term courtship memory was induced as described ^25^. Newly eclosed MB-unc84 males were collected and individually held in an isolation chamber for four to six days. Males were then trained by introducing a single pre-mated female into the isolation chamber for a period of seven hours. After training, males were separated from females and kept in isolation. Flies being used for RNA-seq analysis were collected one-hour after sexual rejection (1h-AR) and 24-hours after rejection 24h-AR. Naïve flies were also collected, and all flies were collected and flash frozen at the same time of day to avoid any gene regulatory effects due to circadian rhythm. Fly heads were isolated from the abdomen, wings, and legs by vortexing followed quickly by separation through a series of sieves. Heads were then stored at −80°C for future processing by INTACT. For each day of courtship conditioning when flies were collected for transcriptome analysis, a subset of naïve and trained males were tested for LTM induction. Statistical significance of courtship suppression was evaluated using a Mann-Whitney *U*-test.

### Isolation of nuclei tagged in a specific cell-type (INTACT)

MB specific transcriptome analysis was accomplished using a described INTACT protocol with several modifications ^33^. Fly heads were then suspended in 1 ml of homogenization buffer (25 mM KCl, 5 mM MgCl2, 20 mM tricine, 0.15 mM spermine, 0.5 mM spermidine, 10 mM β-glycerophosphate, 0.25 mM sucrose, RNAsin Plus RNase Inhibitors (Fisher Scientific: PRN2615), 1X Halt protease inhibitors (Thermo Fisher Scientific: 78430), pH 7.8) and ground with a pestle. To disrupt the cell membrane and release nuclei into solution NP40 was added to the homogenate to an end concentration of 0.3% and the solution was Dounce homogenized 6 times using the tight pestle. The 1 ml nuclear extract was passed through a 40 μm cell strainer and a 50 μl input sample was removed. This input fraction is representative of the whole head, containing both MB-specific GFP nuclei untagged non-MB nuclei. Input fractions were centrifuged to obtain a nuclear pellet which would later be used as a source for whole head RNA sequencing.

Antibody-bound magnetic beads were freshly prepared for each immunopurification by absorbing 1μg of anti-GFP antibody (Invitrogen: G10362) to 60 μl of Protein G Dynabeads (Invitrogen: 10004D) according to the manufacturer’s instructions. To reduce non-specific binding nuclear extracts were pre-cleared by adding 60 μl of beads with no anti-GFP antibody. GFP labeled nuclei were then immunoprecipitated using GFP bound beads for 30 minutes at 4°C with rotation. After washing, these remaining bead-bound nuclei represented the MB-specific fraction that was directly processed for RNA-sequencing.

To investigate the specificity of this protocol a sub-group of MB and WH fractions were incubated with 20mM DRAQ5 (abcam: ab108410) at room temperature for 30 minutes with rotation to label nuclei. Samples were then imaged using a Zeiss AxioImager Z1 and the proportion of GFP positive nuclei to DRAQ5 positive nuclei was determined for three independent biological replicates.

### RNA isolation and RNA-sequencing

RNA was isolated using a PicoPure RNA Isolation Kit (Invitrogen: KIT0204) for both the input and enriched fractions according to the manufacturer instructions. Sequencing libraries were prepared using the Nugen Ovation Drosophila RNA-Seq System 1-16 (Nugen: NU035032) according to instructions. cDNA was then sheared to a target size between 200-300 bp using a Covaris S2 sonicator according to the manufacturer’s protocol. Library size was verified using the Agilent Bioanalyzer High Sensitivity DNA Kit and quantified using a Q-bit fluorometer. Libraries were sequenced on an Illumina NextSeq500 using the high output v2 75 cycle kit to a read length of 75 bp with single-end reads at London Regional Genomics Centre.

### RNA-seq data analysis

Raw sequence reads were trimmed using Prinseq quality trimming to a minimum base quality score of 30 (error probability of 1 in 1,000 base calls) ^63^. Trimmed reads were then aligned to the *D. melanogaster* genome (Ensembl release 88, dm6) using STAR ^64,65^. To ensure mushroom body specificity of MB samples compared to WH samples, we also aligned reads to the *C. elegans* unc-84 gene (NC_003284.9). Only uniquely aligned reads with a maximum of four mismatches were used for downstream analysis. Gene counts were obtained using HTSeq-count using the default union settings to generate genic regions ^66^. To identify differentially expressed genes DESeq2 (R version 3.3.3) was used with cut-offs of q < 0.05, fold change > 1.3 up or down. Genes mapped to the y chromosome were removed from the final DE lists. To identify groups of genes with similar trends of transcriptional regulation in response to courtship conditioning we used the ‘stats’ package in R to perform *k*-means clustering on log2 fold changes ^67,68^.

### GO analysis

Gene ontology (GO) analysis was performed using PANTHER ^69–71^. For GO analysis for biological processes of DE genes between MB and WH samples (**Table ?**) we included all terms with a p < 0.05 (Fisher Exact with FDR multiple test correction). For GO analysis for biological processes of DE genes resulting from courtship conditioning terms were declared significant if they had a p-value of <0.05 (Binomial test with Bonferroni correction). Results are displayed in ‘hierarchical view’ which groups similar terms together under the most enriched term to avoid redundancy^71^. Further functional analysis of the individual genes associated with each enriched term was provided by FlyBase ^72^.

### Network analysis

Interactions network was generated using the GeneMANIA app in Cytoscape 3.4.0 ^73,74^. The network was generated using the following annotated networks: (1) physical interactions – biogrid small scale studies, (2) genetic interactions – biogrid small scale studies, and (3) predicted. No related genes were integrated into the network. Nodes were colour annotated using the Cytoscape enhancedGraphics app ^75^. Each node was annotated based on association with relevant gene ontology terms.

### Data Availability

Supplementary material is available at Figshare. Table S1 contains read alignment and count data. Table S2 contains differential expression analysis results for mushroom body specificity. Table S3 contains gene ontology results for differentially expressed mushroom body enriched or depleted genes. Table S4 contains differential expression analysis results for mushroom body and whole head specific samples during a time course of long-term memory. Table S5 contains the results of k-means clustering of differentially expressed genes during long-term memory formation. Table S6 contains gene ontology results for clusters of deferentially expressed genes identified during long-term memory formation. Gene expression data is available at GEO with the accession number: GSE115718.

## Acknowledgments

This work was funded by a Natural Science and Engineering Research Council of Canada Discovery Grant, the Canada Research Chairs Program, and the Canadian Foundation for Innovation. We thank the Bloomington Drosophila Stock center and the Janelia Research Campus for providing *Drosophila* stocks. Thanks to David Carter and the London Regional Genomics Center for help with sequencing.

